# Gentle, fast and effective crystal soaking by acoustic dispensing

**DOI:** 10.1101/085712

**Authors:** Patrick M Collins, Jia Tsing Ng, Romain Talon, Karolina Nekrosiute, Tobias Krojer, Alice Douangamath, Jose Brandao-Neto, Nathan Wright, Nicholas M Pearce, Frank von Delft

## Abstract

**Synopsis:** A high-throughput method is described for crystal soaking using acoustic droplet ejection, and its effectiveness demonstrated.

**Abstract:** Bright light sources, agile robotics, and fast detectors are continually reducing the time it takes to perform an X-ray diffraction experiment, making high throughput experiments more feasible than ever. But this is also pushing the upstream bottleneck towards sample preparation, even for robust and well characterised crystal systems. Crystal soaking is routinely used to generate protein-ligand complex structures, yet protein crystals are often sensitive to changes in solvent composition, and frequently require gentle or careful stepwise soaking techniques, limiting overall throughput. Here, we describe the use of acoustic droplet ejection for soaking of protein crystals with small molecules, and show that it is both gentle on crystals and allows very high throughput, with 1000 unique soaks easily performed in under 10 minutes. In addition to having very low compound consumption (tens of nanolitres per sample), the positional precision of acoustic droplet ejection enables targeted placement of the compound/solvent away from crystals and towards drop edges, allowing for gradual diffusion of solvent across the drop. This ensures both an improvement in reproducibility of X-ray diffraction and an increased solvent tolerance of the crystals, thus enabling higher effective compound soaking concentrations. We detail the technique here with examples from the protein target JMJD2D, a histone lysine demethylase, having roles in cancer and the focus of active structure based drug design efforts.

## 1. Introduction

Obtaining protein-ligand complexes, the work-horse experiment in structure-based ligand design (SBLD), relies on two methods for achieving the prerequisite crystals: crystal soaking and co-crystallisation (Hassell *et al.*, 2007). Co-crystallisation is achieved by adding the small molecule of interest to the protein prior to setting up a crystallisation experiment, or by simply including it as a component in the crystallisation condition. The potential ligand is free to bind to the protein in solution prior to the formation of a crystal lattice allowing for freedom of potential structural changes. Crystal soaking is the process of taking pre-grown crystals and soaking them with the small molecule of interest. The potential ligand can access the binding sites by diffusing through solvent channels within the crystal lattice, as long as the sites are not involved in crystal packing or otherwise obscured (Danley, 2006).

Crystal soaking tends to be experimentally simpler, since it only requires crystals to be available, and scales well, since a single crystallisation condition can be used to generate a stock of crystals in a known form with higher reliability (Hassell *et al.*, 2007). A frequent obstacle is the low solubility of many compounds solubility in aqueous solutions, requiring organic solvents such as DMSO to be solubilised(Danley, 2006), whereas these solvents also alter the chemistry of the crystal drop and tend to affect the integrity of the crystal. Thus, the basic challenge of crystal soaking is how to introduce the compound to the crystal without destroying the crystal.

A technique that has been gaining utility for protein crystal applications is acoustic droplet ejection, a liquid handling approach that relies on ultrasound pulses focused towards the surface of a liquid, thereby ejecting nanolitre or smaller volume droplets (Ellson *et al.*, 2003). The precision and volume scales of acoustic transfer have enabled new developments in protein crystallography, including: performing small volume crystallisation experiments in crystallisation plates (Wu *et al.*, 2016) or directly on data collection mounts (Yin *et al.*, 2014); transferring pre-formed crystals into mounts (Cuttitta *et al.*, 2015) or directly into the very short pulse of an XFEL beam (Roessler *et al.*, 2016); and preparing high density crystallisation for *in situ* fragment screening (Teplitsky *et al.*, 2015).

An extreme application of crystal soaking is in crystal-based fragment screening. Fragment methods involve screening of a protein target against a library of small molecules, typically under 300 Daltons in size (Congreve *et al.*, 2003). Its advantage is that probability of binding is increased due to the smaller, less complex nature of the molecules (Patel *et al.*, 2014), with chemical elaboration performed on hits to improve potency (Erlanson *et al.*, 2016). Its disadvantage is that the weak binding nature of fragments, a screening method with high sensitivity is required for detection. X-ray crystallography is unrivalled for sensitivity, revealing binding of weakly interacting molecules where other techniques fail (Erlanson *et al.*, 2016), and in this regard is the ideal method for fragment screening (Patel *et al.*, 2014). In addition, the traditional logistical overheads for large scale X-ray experiments are falling away, with the wide-spread availability of fast pixel array detectors and high capacity robotic sample changers (Patel *et al.*, 2014).

We investigated the utility of acoustic droplet ejection for providing a gentle method for crystal soaking at high solvent concentrations, with the requirement of also being rapid and robust for routine large scale crystal soaking as part of the XChem fragment screening user facility at Diamond Light Source.

## 2. Methods

### 2.1. Overview of approach

Our technique for soaking by acoustic dispensing entails transferring compounds in solvent to crystals in sitting drop plates using a Labcyte Echo^®^ 550. Subsequent steps proceed as usual, with drops allowed to soak for a period of time before crystals are harvested, cryo-cooled, and X-ray diffraction data collected.

The Echo operates by moving a transducer below the stationary compound library plate (source plate) and focusing sound pulses at the meniscus of the solution in the requested well, resulting in solvent droplets being ejected upwards (red dots in Figure 1a). The fixed-frequency sound pulse from the transducer in the Echo 550 produces a fixed-sized 2.5 nL droplet, and larger transfer volumes are achieved by dispensing multiple drops of 2.5 nL at a rate of 200 Hz. The inverted sitting drop crystallisation plate (destination plate) is moved above the compound library plate to position the requested target position above the stream of solvent droplets; the relevant wells need to be uncovered during this process.

**Figure 1.**
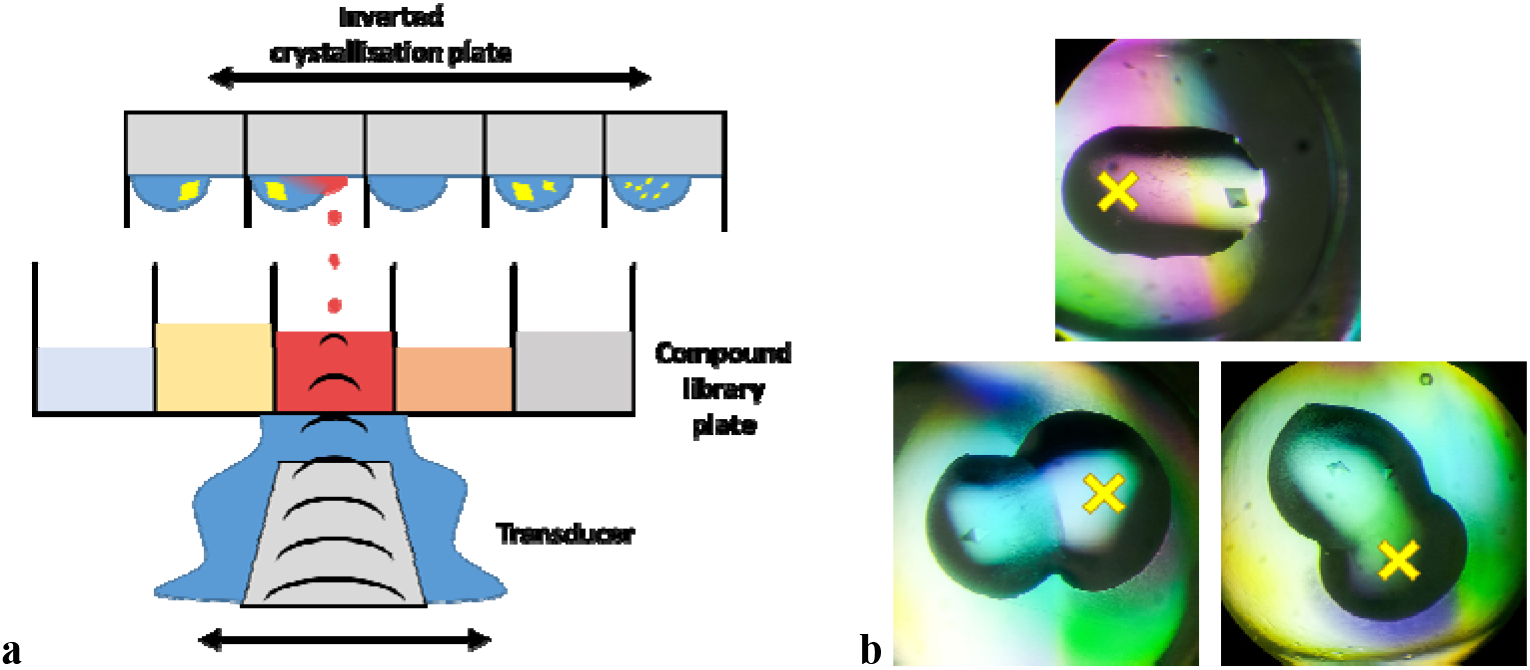
**a** Acoustic droplet ejection for crystal soaking. **b** Protein crystal drops with added compound-containing solvent, acoustically dispensed to the indicated locations (yellow ‘X’) by the offset targeting approach.

In this application, the positional precision of the Echo is relied on to target solvent away from potentially sensitive protein crystals, and towards drop edges (Figure 1b). The compounds used here are dissolved in dimethylsolfoxide (DMSO) at 100 mM concentration, and placed in Labcyte 1536-well source plates. Cryo-protection, when required, is also performed with the Echo by acoustic transfer directly from a 100% solution of ethylene glycol in a 384-well source plate using a mode compatible with viscous solvents (glycerol percentage mode).

### 2.2. Details of acoustic targeting

The Labcyte Plate Reformat software allows for specifying *x y* offset values, in microns, that can be used to specify a target location for acoustic dispensing away from the default centre of the well. In order to build a list of targeted locations, the crystallisation plates were imaged during incubation (Rigaku Minstrel) and images were analysed with TeXRank (Ng *et al.*, 2014). TeXRank uses texture analysis and machine learning methods to rank drops by likelihood of containing a crystal, which greatly facilitates drop selection by ranking the most interesting drops to the beginning of the inspection list (expanded section in Figure 2) presented by the TeXRank visualisation interface. Additionally, TeXRank identifies the centre of each drop-containing lens well from the image, which provides an origin for setting the precise physical location, relative to the centre of the well, to be targeted by the Echo dispensing. The pixel-to-micron scale is also calibrated for a given plate imager.

The TeXRank interface was modified to support targeting by a single mouse-click. Clicking on a specific point of the image (yellow ‘X’, Figure 2): a) registers the drop for inclusion in the experiment (contains a suitable crystal), b) records the target location for acoustic dispensing, an *x y* coordinate offset from the origin (well centre) in microns, and c) brings up the next well in the ranked list. Thus, a crystallographer can very rapidly (in minutes) build up the whole list of very precise dispensing target locations. This list, consisting of plate name (usually a barcode), well location, and *x y* offset coordinates is exported from TeXRank. The file format output by TeXRank is detailed in section S1.

**Figure 2.**
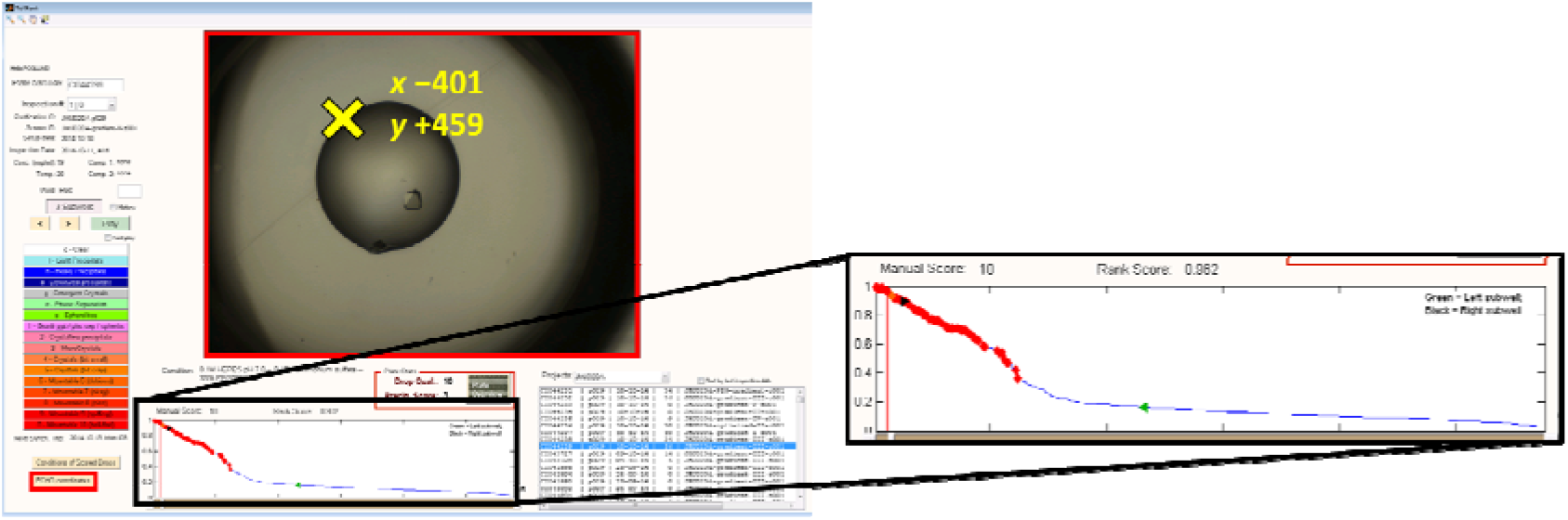
The TeXRank interface showing a crystallisation drop containing a single JMJD2D crystal. Clicking a location records the acoustic dispensing target (yellow ‘X’ and *x y* coordinates have been added for clarity). Expanded section shows the ranked plot of crystal images.

### 2.3. Configuring the Echo

The sitting drop crystallisation plates used were SWISSCI 3-drop plates (Ng *et al.*, 2016). The Echo is compatible with arbitrary destination plates which can be configured within the software by creation of a labware definition for the plate. The SWISSCI 3-drop plates (Figure 3) have 96 positions with 4 subwells (3 crystallisation wells, and 1 reservoir well) per position. The labware definition for such a plate can be created and used within the Labcyte Array Maker software; however this software is not compatible with the targeting offset values, nor does it have the ability to import transfer lists created outside the software. Instead, the plate was defined in a 384-well format (red numbering in Figure 3) and used with the Labcyte Plate Reformat software, which does allow for targeting offset values and importing of transfer lists. To address the technical complication that wells are non-uniformly distributed horizontally, with subwell ‘d’ (Figure 3) positioned 700 microns off-centre, an additional offset correction of 700 microns is applied to these positions (even-numbered columns in 384-well format). The Echo labware definition for the SWISSCI 3-drop plate definition in 384-well format is available for download from the Diamond XChem website. Further fine-tuning of accuracy was performed by iterative dispensing and adjustment to the plate definition, and mechanical calibration of the destination plate carrier.

**Figure 3.**
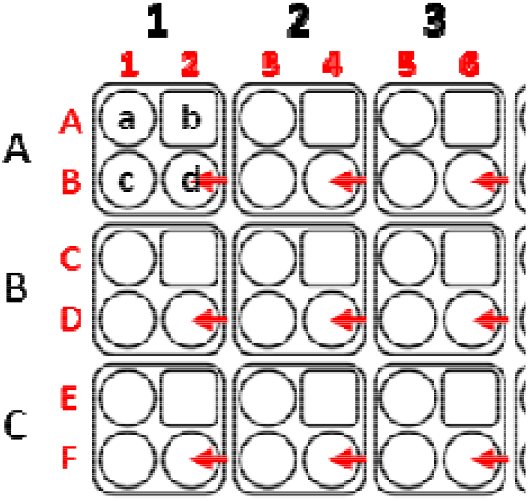
Top left corner of a SWISSCI 3 lens crystallisation plate. The normal definition of the plate has 96 locations, with four subwells per location (black text). We define the plate in a 384-well format (red text) and apply an offset correction to the even-numbered columns (red arrows), which are not positioned centrally between adjacent subwells.

Lists of acoustic dispensing targets from TeXRank can be matched to a list of compounds with any spreadsheet tool. In this work, transfer volumes were calculated by to achieve given final compound or solvent concentrations assuming the crystallisation drop volume was that initially dispensed. The file format required for upload to the Echo is detailed in section S2.

### 2.4. Crystallisation, data collection, processing, and refinement

JMJD2D (KDM4D) was expressed and purified as previously described (Bavetsias *et al.*, 2016). Crystals were grown in SWISSCI 3 lens crystallisation sitting drop plates at 20 °C by mixing 50–100 nL of 11 mg/mL protein in a 1:1 ratio with 50–100 nL of reservoir solution containing 0.1 M HEPES pH 7.0, 0.15 M ammonium sulfate, and 26–37 % (w/v) PEG3350, and placed over 20 μL of reservoir solution. Crystals appeared in 1–3 days. Crystal soaking was performed with acoustic transfer using a Labcyte Echo 550, with compounds/solvent targeted away from crystals and towards drop edges. Ethylene glycol was added for cryoprotection to 20%, calculated from the initial drop volume, and added to a targeting location using the echo. JMJD2DA crystals diffracted to 1.3–1.6 Å resolution, in space group *P*4_3_2_1_2 with typical unit cell dimensions of a=71.5 Å, c=150 Å with one JMJD2D molecule in the asymmetric unit.

X-ray diffraction data were collected at Diamond Light Source beamline I04-1 and processed through the Diamond autoprocessing pipeline, which utilises *xia2* (Winter, 2010), *DIALS* (Waterman *et al.*, 2016), *XDS* (Kabsch, 2010), *pointless* (Evans, 2006), and *CCP4* (Winn *et al.*, 2011). Electron density maps were generated using *XChemExplorer* (Krojer, 2016 reference in this issue) via *DIMPLE* (Wojdyr *et al.*, 2013). Ligand restraints were generated with *ACEDRG* and ligand binding was detected with *PanDDA* (Pearce *et al.*, 2016), with ligands built into *PanDDA* event maps. Iterative refinement and manual model correction was performed using *REFMAC* (Murshudov *et al.*, 2011) and *COOT* (Emsley *et al.*, 2010), respectively.

## 3. Results and discussion

### 3.1 Acoustic dispensing is effective for soaking

In order to confirm that acoustic dispensing could be used effectively for obtaining protein-ligand complexes, we performed soaking experiments using crystals of the histone demethylase JMJD2D and a known binder reported as part of a structure-based design effort targeting the histone lysine demethylase (KDM) family (compound 30a from Bavetsias et al., 2016, named KDOAM16 here). A number of acoustic transfers were performed to different crystal drops, with transfer volumes ranging from 5 to 50 nL (directly from a 100 mM DMSO stock). Electron density maps from all experiments clearly revealed strong positive difference density for the ligand in the binding site (Figure 4). Refinement of the ligand showed that the binding pose and protein-ligand interactions of KDOAM16 were identical to those previously reported (PDB ID: 5F5A). This formed the basis of further experiments to identify optimal soaking parameters and experimental procedures, with a view to deploying the technique as a robust and central component of the XChem fragment screening facility at Diamond Light Source.

**Figure 4.**
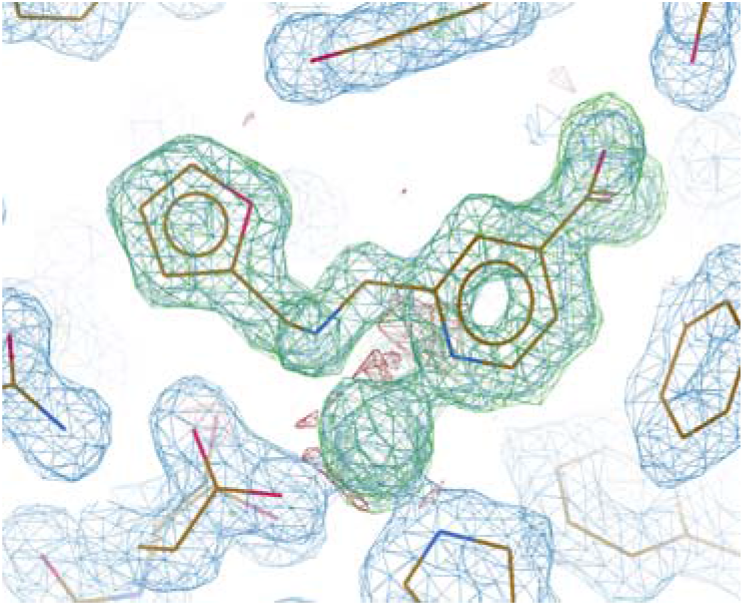
Electron density maps showing bound KDOAM16 (σA-weighted 2m*F*_o_×D*F*_c_: blue, 1.4 σ, and m*F*_o_×D*F*_c_: red/green, ±3 σ 1.5 Å resolution) calculated from refinement prior to including the ligand in the model.

### 3.2 Dispense volumes and tolerated concentrations

Although the Echo has high precision (<8 % CV) for small volume transfers, (Ellson *et al.*, 2005), an accurate estimate of the final concentration of solvent or compound after acoustic transfer requires the volume of the crystallisation drop to be known. Instead, only the initial drop volume prior to vapour diffusion is available. In a vapour diffusion experiment, the drop volume of a 1:1 protein:reservoir solution will typically reduce to approximately half the original volume, although the exact end volume depends on presence of solutes in the protein component, which compete with the diffusion processes (Luft & DeTitta, 2008). This is especially true when PEG solutions are used in the reservoir, since PEGs do not reduce vapour pressure of water as effectively compared to salts (Luft & DeTitta, 2008). Establishing such details for many different crystallization systems is not practicable, and therefore no attempt was made here to estimate final drop volumes.

Instead we perform a solvent tolerance screen on new conditions to determine the exact amounts of solvent tolerated by the crystal system under the acoustic dispensing conditions (discussed further in section 3.7). It should nevertheless be highlighted that the final concentration (solvent percentage or compound concentration) reported here are calculated based on the initial drop volume, and are likely underestimates of the true final concentration by up to half, in the case of 1:1 protein:reservoir drops.

### 3.3 Exploiting positional precision

The ability of the Echo to dispense solvent to arbitrarily requested locations within a subwell, or to dispense with a complex pattern in 2.5 nL units (Figure 5), opens up possibilities for crystal soaking experiments that are not possible to perform by hand. On the other hand, compound is dispensed directly from 100 % stock solutions and sudden additions of solvent-altering components is in general stressful for sensitive protein crystals. We therefore investigated how the positional precision of acoustic transfer could be exploited to ensure soaking was not only rapid but also sufficiently gentle.

**Figure 5.**
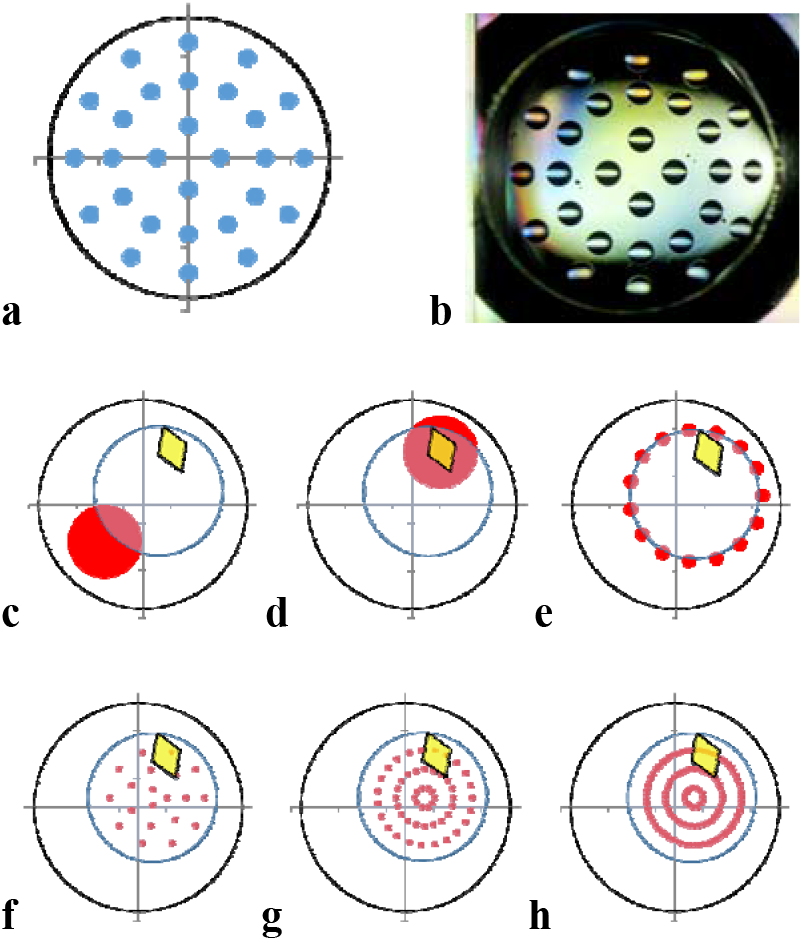
Positional precision of acoustic transfer. **a** The requested pattern of dispensing, and **b** the resulting 2.5 nL drop pattern within a subwell of a sitting drop plate. Other dispensing patterns investigated were **c**) an offset location away from the crystal, **d** crystal targeting, **e** a ring pattern around the drop edge, or multiple ring patterns across the drop, increasing in density with **f** 20 %, **g** 40%, or **h** 60 % solvent concentrations.

**Figure 6.**
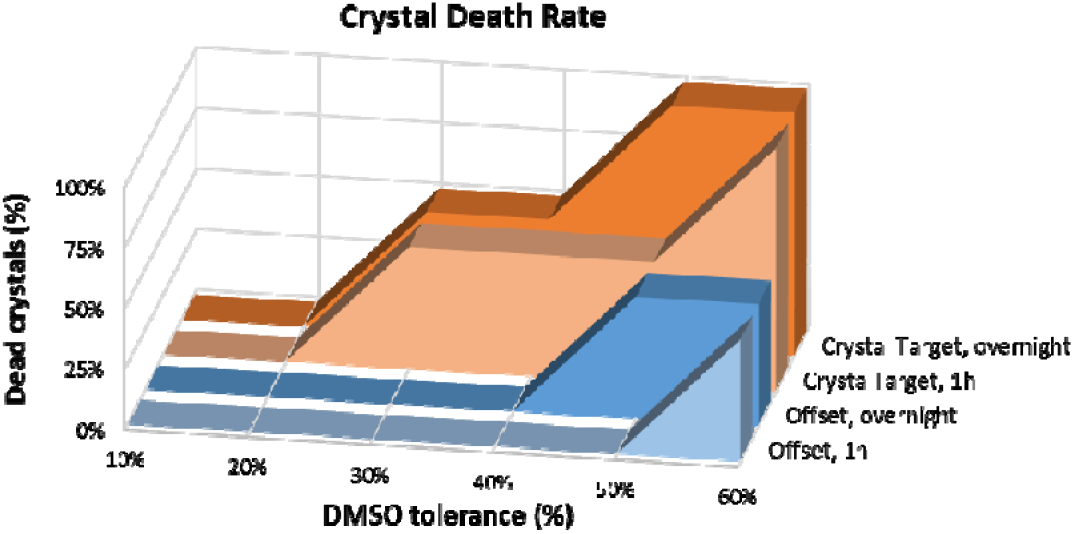
Death rate of JMJD2D crystals from soaking in DMSO after acoustic transfer targeted at the crystal (orange), or targeted away from the crystal (blue). Crystals were soaked for 1 hour (lighter colour), or overnight (darker colour).

All of the dispensing patterns in Figure 5 (c–h) were investigated with JMJD2D crystals in duplicate with concentrations ranging from 10–60 % solvent, and with 1 hour and overnight soaking times (96 soaking experiments), with crystal survival measured by X-ray diffraction. It was found that crystals tolerate twice the solvent concentration when targeted away from the crystal with the offset approach compared to when the crystal was directly targeted. Targeted crystals tolerated 20 % solvent, whereas offset targeting provided 40 % solvent tolerance (Figure 6**Error! Reference source not found.**). Intermediate survival rates were observed for the different ring patterns, with crystals tolerating up to 30 % DMSO (data not shown).

### 3.4. Diffusion

Typical crystal soaking experiments usually involve preparing the compound of interest in a crystal compatible solution, usually reservoir solution with additional solvent such as DMSO, after which either the solution is transferred directly to the crystallisation drop, or the crystal is moved to the solution. This exposes the crystal to sudden changes from its native solution, which can damage the crystal through osmotic shock (Lopez-Jaramillo *et al.*, 2002). Some crystals are far more tolerant to this form of treatment than others, but for those that are not, careful stepwise procedures can overcome this, enabling higher solvent and compound concentrations to be introduced gradually to the crystal (Hassell *et al.*, 2007).

This provides the most likely explanation for why crystals tolerate the very high solvent concentrations generate by the acoustic offset targeting described here. Gradual diffusion of solvent/solute across the drop allows the crystal a significantly longer equilibration time (Figure 7a). A similar observation has been reported when compounds are delivered via a laser-generated aperture (Zander *et al.*, 2016). We observe that it takes 2–5 minutes for coloured compounds or dyes to diffuse across the drop and reach an equilibrium after acoustic transfer to the edge of a crystallisation drop. In contrast, crystals that are plunged into a drop containing a new solvent, or are flooded by addition of solvent, will experience equilibration within milliseconds or seconds at best, 2-5 orders of magnitude faster than gradual diffusion from offset targeting.

When a skin is present on the crystallisation drop it acts as a membrane partition between the crystal drop and the transferred solution, but still allows gradual diffusion of solutes (Figure 7b). The process is however slower only by an order of magnitude, minutes to hours as judged by colour equilibration, depending on the age of the drop and thickness of the skin. Disruption of skin with a loop or micro tool results in influx of compound over a number of minutes as observed in the absence of a skin. Overall, a skin does not prevent acoustic crystal soaking, however appropriate soaking times will need to be established.

**Figure 7.**
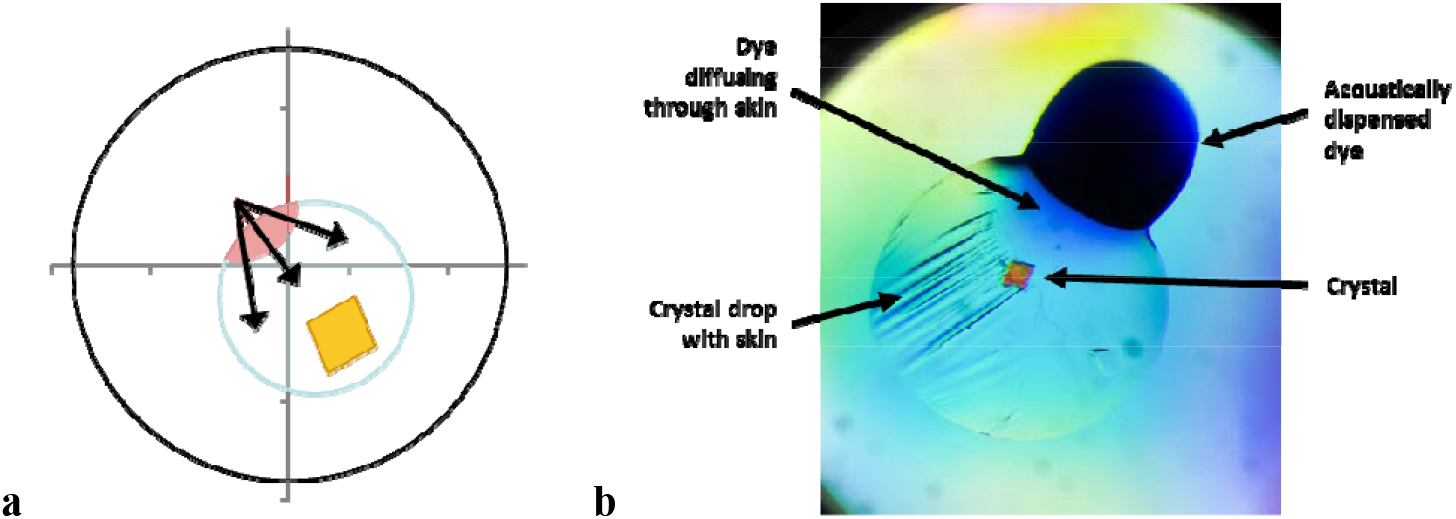
**a** Schematic illustration of diffusion (black arrows) from a drop soaked with the offset targeting approach. **b** Reduced diffusion of methylene blue dye through a crystallisation drop that has a skin.

Offset targeting is now the default protocol for the XChem platform, for three reasons. Firstly, it allows very high solvent concentrations to be dispensed, and it is reasonable to assume a the correspondingly high compound concentration is important for ensuring the weakly binding fragments bind with sufficient occupancy to ensure detection in electron density maps Secondly, offset targeting is simple to perform, since a single click in TeXRank defines the target. Thirdly, overall dispensing speed is considerably faster for a single target compared to the more complex partners, which require many additional stage movements within the Echo (section 3.5).

The same targets used for compound dispensing can also be used for adding cryoprotecting solutions to the drop prior to harvesting. Ethylene glycol is a convenient cryoprotectant since acoustic transfer can be performed directly from a 100 % stock solution. The higher viscosity of glycerol requires a 50 % diluted stock solution for successful acoustic transfer, and thus requires larger volumes to be added to achieve the required cryoprotecting concentration. However, for the routine fragment screening at Diamond, we, like others (Pellegrini *et al.*, 2011, Zander *et al.*, 2016), have observed that by matching mounting loop to crystal size and limiting excessive solvent surrounding the crystal, cryoprotection is often not required (section 3.7).

### 3.5. Speed and throughput

Acoustic dispensing is rapid for small nanolitre scale volumes once the experiment has been designed and a transfer list constructed, as described in sections 2.2 and 2.3. The fixed rate of droplet ejection, 2.5 nL at 200 Hz, indicates a fluid transfer rate for 500 nL/s; however, for the large numbers of <100 nL transfers typical for fragment screening, the actual throughput is limited by stage movements rather than fluid transfer. For example, 1000 × 25 nL transfers take 7 minutes, but only 8.5 minutes for the same number of 100 nL transfers (62 nL/s and 196 nL/s, respectively). Therefore, significant changes in transfer volumes have only a marginal impact of total transfer times at these scales, which correspond to the usual crystallisation drop volumes, typical for robotically prepared crystallisation experiments (100–200 nL).

In practice, therefore, the transfer of 1000 unique compounds to 1000 unique crystallisation drop locations can be performed in under 10 minutes. This includes the time taken for stage movements within the Echo, and unsealing and resealing 4–5 crystallisation plates full of crystals.

For the typical crystallisation drop sizes cited above, the 1–2 minute timeframe for acoustic dispensing per plate is short enough that the plate seal can be completely removed during transfer without evaporation affecting drops or crystal integrity (results not shown). The internal chamber of the Echo is also somewhat humidified, as the sound waves are coupled from the transducer to the bottom of the source plate by flowing water, and presumably this helps slow down evaporation.

For smaller drop volumes or volatile crystallisation components, the experiment can be broken into batches by exposing smaller sections of the plate at a time; the only drawback is that the overall experiment takes more time. It has also been reported that the use of a plate lid containing small apertures that allow acoustic transfer but minimises disruption of the vapour environment around the drop, significantly reduces evaporation and improves X-ray diffraction consistency (Zipper *et al.*, 2014).

### 3.6. Time and concentration

In order to investigate the effectiveness of acoustic transfer for crystal soaking and ligand binding, a number of experiments were performed with known binders of varying affinities under different conditions. Two molecules identified from fragment screening of JMJD2D (unpublished, models available from http://www.thesgc.org/fragment-screening) were selected and categorised as a <u>medium</u> binder and a <u>weak</u> binder, based on the signal from previously observed electron density maps. Also included was compound KDOAM16 (Bavetsias *et al.*, 2016) (section 3.1), which was designated the <u>strong</u> binder. Actual binding affinities for these molecules with JMJD2D have not yet been measured.

Figure 8a shows the detection threshold of the three molecules as a function of concentration. The <u>weak</u> binder was only observed at 30 mM concentration, and then only for one out of the two duplicates, while the <u>medium</u> binder was not detected below 20 mM. The <u>tight</u> binder was detected for both duplicates at the minimum concentration tested of 1.2 mM. This corresponded to a single 2.5 nL acoustic droplet transferred to the crystallisation drop, and shows that for tight binding ligands, use of acoustic transfer is extremely effective, requiring only very limited amounts of compound. Two 2.5 nL transfers directly from the intact 100 mM sock (116 ng total compound) were sufficient to obtain two separate protein-ligand complex structures (Figure 9). The results for the medium and weak binding ligands highlight the importance of compound concentration in order to detect weak binding ligands. Since the crystals have a ceiling on solvent tolerance, the next best way to increase compound concentration is to increase the concentration of the stock solution, though in practice, compound solubility will dictate the effective concertation that can be achieved, either in the stock solution or after addition the crystallisation drop.

**Figure 8.**
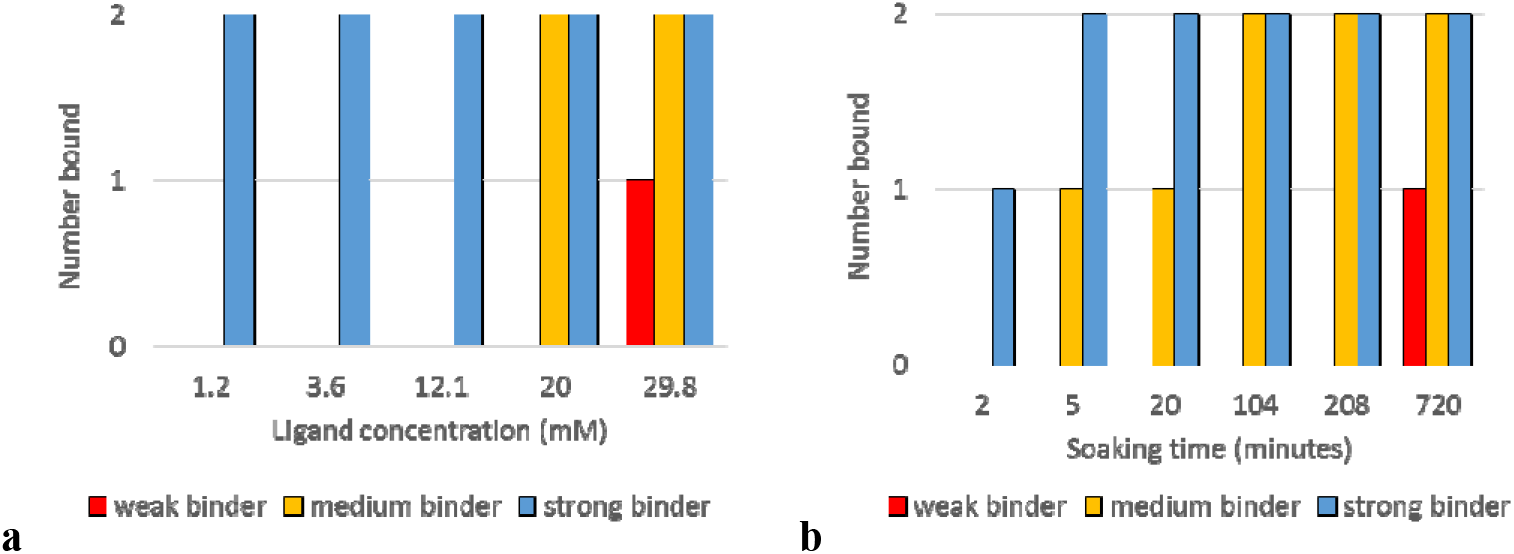
The detection of strong, medium, and weak binding ligands as a function of **a** concentration, or **b** time, after crystal soaking using acoustic transfer (from 100 mM stock). The concentration series in **a** were soaked for a fixed time of 4 hours, while the time series in **b** were soaked at a fixed concentration of 20 mM. Soaks were performed in duplicate for each condition leading to 30 X-ray diffraction data sets for **a** and 36 datasets for **b**.

The detection of binding for the three molecules from soaking for different lengths of time after acoustic transfer (20 mM soaks) is shown in Figure 8b. For the <u>strong</u> binder, only one of the duplicates was observed when the crystals were mounted 2 minutes after transfer. Care was taken when mounting to pick the crystal directly out of the drop without excessive mixing of solvent within the drop, in order to isolate the effect of compound diffusion across the drop; nevertheless, some mixing will inevitably have occurred. For the <u>medium</u> binder, 2 minutes was insufficient for the ligand to be detected, while at 5 minutes, one of the duplicates was detected (Figure 9). The <u>weak</u> binder was only detected after an overnight soak at 20 mM.

**Figure 9.**
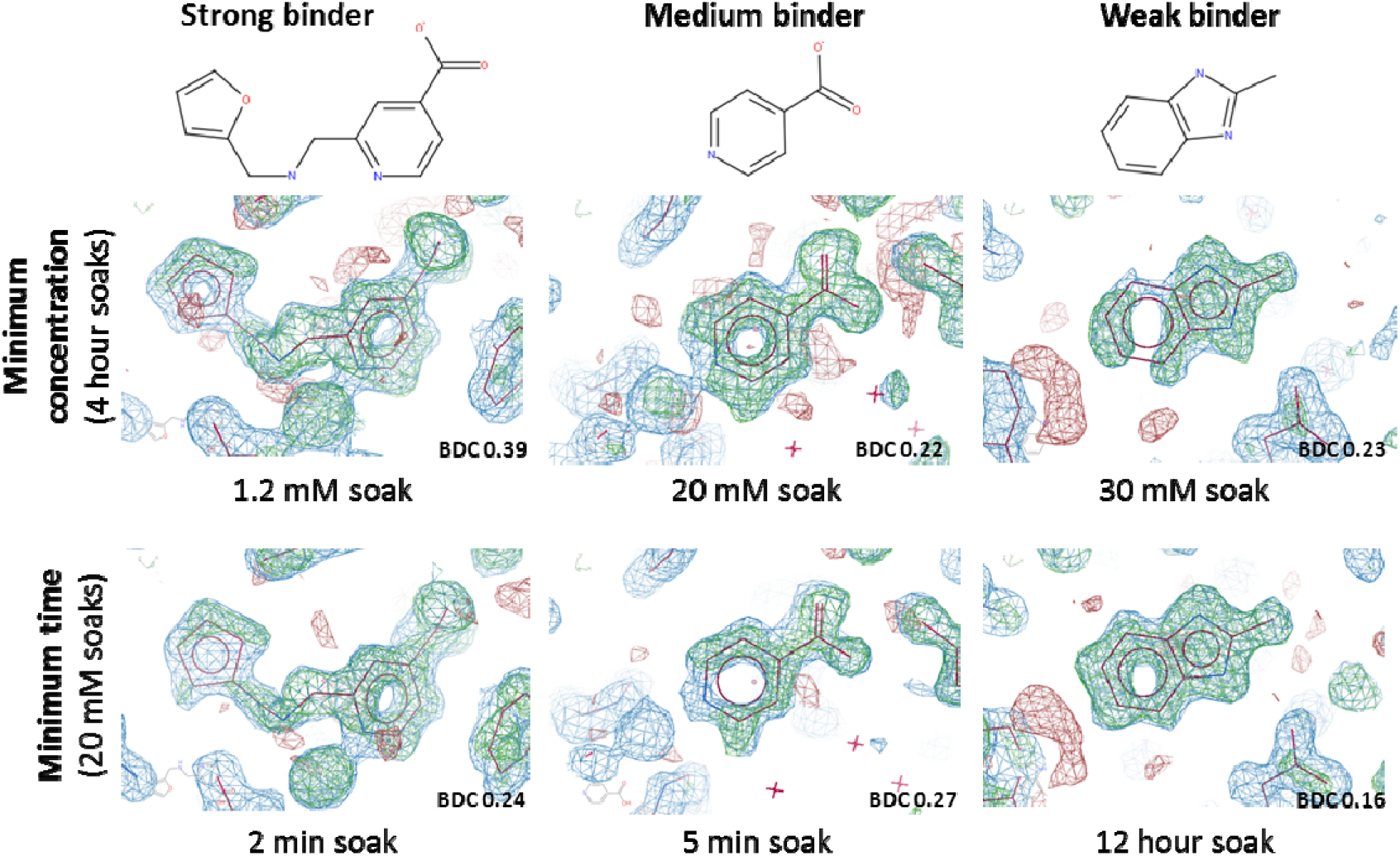
Electron density maps (PanDDA maps: event maps, blue, 2 σ, Z-maps: green/red, ±3, 1.3–1.4 Å resolution) from the minimum experimental conditions (time or concentration series) required to detect ligand binding. The PanDDA reported background density correction (BDC) values are shown (Pearce *et al.*, 2016).

### 3.7. The importance of control experiments

As discussed above, the dynamics of adding small amounts of 100 % stock solutions directly to a crystallisation drop for crystal soaking are different compared to manually transferring a crystal to a new drop, or flooding it with a pre-prepared solution. From many subsequent experiments (not reported here) we conclude that previous knowledge of solvent tolerance of a crystal system tends to be unrelated to that observable by acoustic dispensing with offset targeting, which typically permits dramatically higher solvent concentrations.

The strong implication is that thorough advance control experiments are essential for establishing a maximally effective offset targeting protocol. A standard solvent tolerance screen has been implemented, that involves testing X-ray diffraction from crystals soaked at 5–40 % final DMSO concentration, at 3 time points (1 hour, 3 hours, and overnight) in duplicate, and including untreated control crystals (30–36 crystals in total). From over 18,000 crystal soaks across 16 recent protein targets, the DMSO solvent tolerance was found to range between 10–40 % (23 % on average, of course, the actual concentrations are likely underestimated by as much as half, as discussed in section 3.2), with soaking times of 4–6 hours (Figure 10). Similarly, 11 of the 16 targets have not required further cryoprotection. Finally, the real-life validity of the parameters are confirmed by performing a significant number of soaking experiments (10-100) to ensure diffraction is indeed consistently retained.

**Figure 10.**
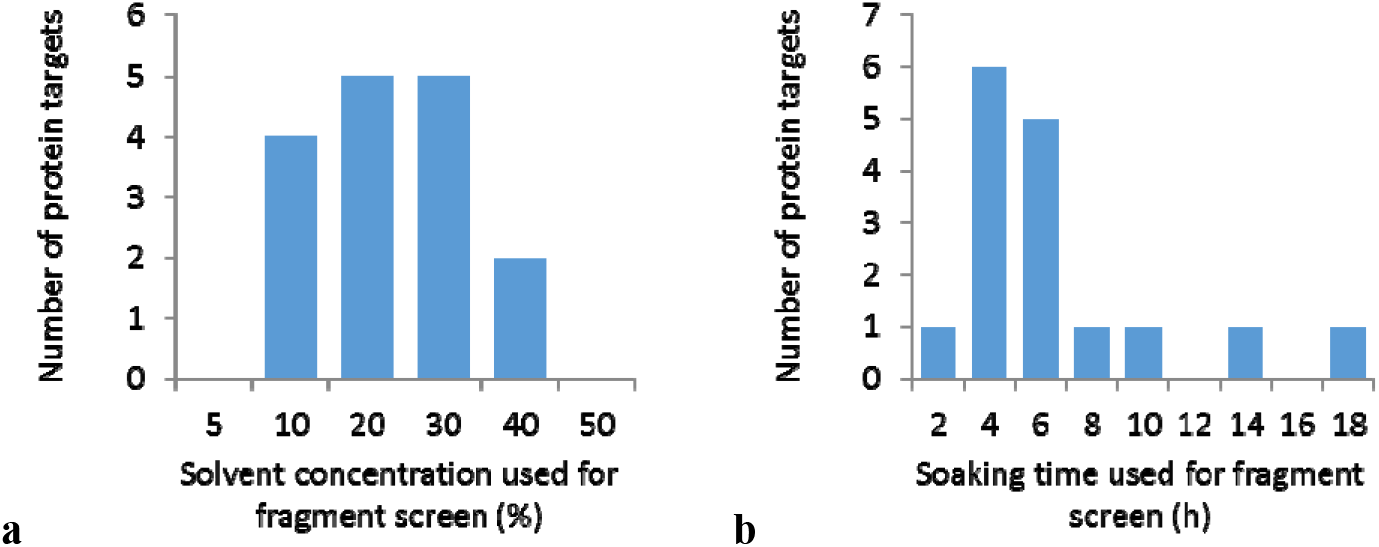
Working experimental parameters for **a** DMSO solvent concentration and **b** soaking time from large scale fragment screening of 16 protein targets, totalling over 18,000 crystal soaks.

## 4. Conclusions

We have developed a method for soaking of protein crystals that is both gentle and rapid. By using the precision of acoustic dispensing to target the transfer of solvent and compounds away from sensitive protein crystals, an increase in solvent tolerance and X-ray diffraction reproducibility can be achieved, even at a rate of 100 crystals per minute. Use of control experiments to empirically determine optimal experimental conditions for each crystal system has enabled fragment screening on multiple diverse protein targets in the XChem facility at Diamond.

## Acknowledgements

We would like to thank the Diamond GDA team, Scientific software team, and Experimental Hall coordinators for ensuring that the data kept flowing. We would also like to thank Celine Be, Novartis, for early discussions and ideas, and Aleksandra Szykowska, SGC, for purified protein. The SGC is a registered charity (No. 1097737) that receives funds from AbbVie, Bayer, Boehringer Ingelheim, the Canada Foundation for Innovation, the Canadian Institutes for Health Research, Genome Canada, GlaxoSmithKline, Janssen, Lilly Canada, the Novartis Research Foundation, the Ontario Ministry of Economic Development and Innovation, Pfizer, Takeda and the Wellcome Trust (092809/Z/10/Z).

## Supporting information

### S1. TeXRank output csv file format

The output format for TeXRank is a csv file with 4 columns. For a SWISSCI 3 drop plate, the file contains all 288 wells (288 lines in the file) (no header), with those selected for targeting containing the *x y* coordinates. Column 1: Well Number, Column 2: *x* offset value, Column 3: *y* offset value, Column 4: a score (default = 6)

For example, the first 5 lines of a TeXRank output file where 3 drops have been targeted, and 2 wells are not targeted.

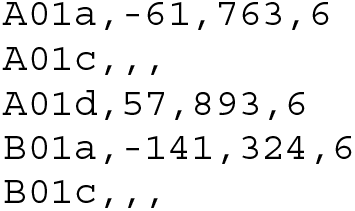

### S2. Echo region definition input file format

The region definition input file for the Labcyte Plate Reformat software is a csv file with 6 columns, and includes a header row. Column 1: Palate name, Column 2: Source well, Column 3: Destination well, Column 4: Transfer Volume, Column 5: Destination Well X offset, Column 6: Destination Well Y Offset.

For example, the first 6 lines of an Echo region definition csv file:

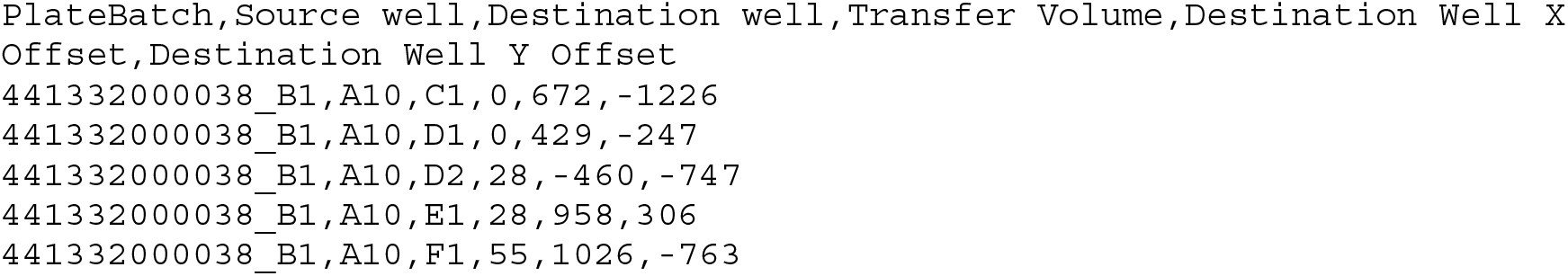

## References

Bavetsias, V., Lanigan, R. M., Ruda, G. F., Atrash, B., McLaughlin, M. G., Tumber, A., Mok, N. Y., Le Bihan, Y. V., Dempster, S., Boxall, K. J., Jeganathan, F., Hatch, S. B., Savitsky, P., Velupillai, S., Krojer, T., England, K. S., Sejberg, J., Thai, C., Donovan, A., Pal, A., Scozzafava, G., Bennett, J. M., Kawamura, A., Johansson, C., Szykowska, A., Gileadi, C., Burgess-Brown, N. A., von Delft, F., Oppermann, U., Walters, Z., Shipley, J., Raynaud, F. I., Westaway, S. M., Prinjha, R. K., Fedorov, O., Burke, R., Schofield, C. J., Westwood, I. M., Bountra, C., Muller, S., van Montfort, R. L., Brennan, P. E. & Blagg, J. (2016). Journal of medicinal chemistry 59, 1388–1409.

Congreve, M., Carr, R., Murray, C. & Jhoti, H. (2003). Drug discovery today 8, 876–877.

Cuttitta, C. M., Ericson, D. L., Scalia, A., Roessler, C. G., Teplitsky, E., Joshi, K., Campos, O., Agarwal, R., Allaire, M., Orville, A. M., Sweet, R. M. & Soares, A. S. (2015). Acta Crystallographica Section D 71, 94–103.

Danley, D. E. (2006). Acta crystallographica. Section D, Biological crystallography 62, 569–575.

Ellson, R., Mutz, M., Browning, B., Lee, L., Miller, M. F. & Papen, R. (2003). Journal of the Association for Laboratory Automation 8, 29–34.

Ellson, R., Stearns, R., Mutz, M., Brown, C., Browning, B., Harris, D., Qureshi, S., Shieh, J. & Wold, D. (2005). Combinatorial chemistry & high throughput screening 8, 489–498.

Emsley, P., Lohkamp, B., Scott, W. G. & Cowtan, K. (2010). Acta crystallographica. Section D, Biological crystallography 66, 486–501.

Erlanson, D. A., Fesik, S. W., Hubbard, R. E., Jahnke, W. & Jhoti, H. (2016). Nat Rev Drug Discov 15, 605–619.

Evans, P. (2006). Acta Crystallographica Section D 62, 72–82.

Hassell, A. M., An, G., Bledsoe, R. K., Bynum, J. M., Carter, H. L., 3rd, Deng, S. J., Gampe, R. T., Grisard, T. E., Madauss, K. P., Nolte, R. T., Rocque, W. J., Wang, L., Weaver, K. L., Williams, S. P., Wisely, G. B., Xu, R. & Shewchuk, L. M. (2007). Acta crystallographica. Section D, Biological crystallography 63, 72–79.

Kabsch, W. (2010). Acta crystallographica. Section D, Biological crystallography 66, 125–132.

Lopez-Jaramillo, F. J., Moraleda, A. B., Gonzalez-Ramirez, L. A., Carazo, A. & Garcia-Ruiz, J. M. (2002). Acta crystallographica. Section D, Biological crystallography 58, 209–214.

Luft, J. R. & DeTitta, G. T. (2008). Protein crystallization, Second ed., edited by T. M. Bergfors, pp. 11–46. La Jolla, Calif: International University Line.

Murshudov, G. N., Skubak, P., Lebedev, A. A., Pannu, N. S., Steiner, R. A., Nicholls, R. A., Winn, M. D., Long, F. & Vagin, A. A. (2011). Acta crystallographica. Section D, Biological crystallography 67, 355–367.

Ng, J. T., Dekker, C., Kroemer, M., Osborne, M. & von Delft, F. (2014). Acta crystallographica. Section D, Biological crystallography 70, 2702–2718.

Ng, J. T., Dekker, C., Reardon, P. & von Delft, F. (2016). Acta Crystallogr D Struct Biol 72, 224–235.

Patel, D., Bauman, J. D. & Arnold, E. (2014). Progress in biophysics and molecular biology 116, 92- 100.

Pearce, N., Bradley, A. R., Collins, P., Krojer, T., Nowak, R., Talon, R., Marsden, B. D., Kelm, S., Shi, J., Deane, C. & von Delft, F. (2016). bioRxiv.

Pellegrini, E., Piano, D. & Bowler, M. W. (2011). Acta Crystallographica Section D 67, 902–906.

Roessler, C. G., Agarwal, R., Allaire, M., Alonso-Mori, R., Andi, B., Bachega, J. F., Bommer, M., Brewster, A. S., Browne, M. C., Chatterjee, R., Cho, E., Cohen, A. E., Cowan, M., Datwani, S., Davidson, V. L., Defever, J., Eaton, B., Ellson, R., Feng, Y., Ghislain, L. P., Glownia, J. M., Han, G., Hattne, J., Hellmich, J., Heroux, A., Ibrahim, M., Kern, J., Kuczewski, A., Lemke, H. T., Liu, P., Majlof, L., McClintock, W. M., Myers, S., Nelsen, S., Olechno, J., Orville, A. M., Sauter, N. K., Soares, A. S., Soltis, S. M., Song, H., Stearns, R. G., Tran, R., Tsai, Y., Uervirojnangkoorn, M., Wilmot, C. M., Yachandra, V., Yano, J., Yukl, E. T., Zhu, D. & Zouni, A. (2016). Structure 24, 631- 640.

Teplitsky, E., Joshi, K., Ericson, D. L., Scalia, A., Mullen, J. D., Sweet, R. M. & Soares, A. S. (2015). Journal of Structural Biology 191, 49–58.

Waterman, D. G., Winter, G., Gildea, R. J., Parkhurst, J. M., Brewster, A. S., Sauter, N. K. & Evans, G. (2016). Acta Crystallographica Section D 72, 558–575.

Winn, M. D., Ballard, C. C., Cowtan, K. D., Dodson, E. J., Emsley, P., Evans, P. R., Keegan, R. M., Krissinel, E. B., Leslie, A. G., McCoy, A., McNicholas, S. J., Murshudov, G. N., Pannu, N. S., Potterton, E. A., Powell, H. R., Read, R. J., Vagin, A. & Wilson, K. S. (2011). Acta crystallographica. Section D, Biological crystallography 67, 235–242.

Winter, G. (2010). Journal of Applied Crystallography 43, 186–190.

Wojdyr, M., Keegan, R., Winter, G. & Ashton, A. (2013). Acta Crystallographica Section A 69, s299.

Wu, P., Noland, C., Ultsch, M., Edwards, B., Harris, D., Mayer, R. & Harris, S. F. (2016). J Lab Autom 21, 97–106.

Yin, X., Scalia, A., Leroy, L., Cuttitta, C. M., Polizzo, G. M., Ericson, D. L., Roessler, C. G., Campos, O., Ma, M. Y., Agarwal, R., Jackimowicz, R., Allaire, M., Orville, A. M., Sweet, R. M. & Soares, A. S. (2014). Acta crystallographica. Section D, Biological crystallography 70, 1177–1189.

Zander, U., Hoffmann, G., Cornaciu, I., Marquette, J.-P., Papp, G., Landret, C., Seroul, G., Sinoir, J., Rower, M., Felisaz, F., Rodriguez-Puente, S., Mariaule, V., Murphy, P., Mathieu, M., Cipriani, F. & Marquez, J. A. (2016). Acta Crystallographica Section D 72, 454–466.

Zipper, L. E., Aristide, X., Bishop, D. P., Joshi, I., Kharzeev, J., Patel, K. B., Santiago, B. M., Joshi, K., Dorsinvil, K., Sweet, R. M. & Soares, A. S. (2014). Acta Crystallographica Section F 70, 1707- 1713.

